# TFBSFootprinter: a multiomics tool for prediction of transcription factor binding sites in vertebrate species

**DOI:** 10.1101/2025.01.09.632105

**Authors:** Harlan R. Barker, Seppo Parkkila, Martti E.E. Tolvanen

## Abstract

**Background:** Transcription factor (TF) proteins play a critical role in the regulation of eukaryotic gene expression via sequence-specific binding to genomic locations known as transcription factor binding sites (TFBSs). Accurate prediction of TFBSs is essential for understanding gene regulation, disease mechanisms, and drug discovery. These studies are therefore relevant not only in humans but also in model organisms and domesticated and wild animals. However, current tools for the automatic analysis of TFBSs in gene promoter regions are limited in their usability across multiple species. To our knowledge, no tools currently exist that allow for automatic analysis of TFBSs in gene promoter regions for many species.

**Methodology and Findings:** The TFBSFootprinter tool combines multiomic transcription-relevant data for more accurate prediction of functional TFBSs in 124 vertebrate species. In humans, this includes vertebrate sequence conservation (GERP), proximity to transcription start sites (FANTOM5), correlation of expression between target genes and TFs predicted to bind promoters (FANTOM5), overlap with ChIP-Seq TF metaclusters (GTRD), overlap with ATAC-Seq peaks (ENCODE), eQTLs (GTEx), and the observed/expected CpG ratio (Ensembl).

TFBSFootprinter analyses are based on the Ensembl transcript ID for simplicity of use and require minimal setup steps. Benchmarking of the TFBSFootprinter on a manually curated and experimentally verified dataset of TFBSs produced superior results when using all multiomic data (average area under the receiver operating characteristic curve, 0.881), compared with DeepBind (0.798), FIMO (0.817) and traditional PWM (0.854). The results were further improved by selecting the best overall combination of multiomic data (0.910). Additionally, we determined combinations of multiomic data that provide the best model of binding for each TF. TFBSFootprinter is available as Conda and Python packages.

## Introduction

Transcription factor (TF) proteins play a critical role in the regulation of eukaryotic gene expression by sequence-specific binding to short stretches of DNA (6–24 bp) known as transcription factor binding sites (TFBSs) [1] which can comprise larger genomic locations known as cis-regulatory elements (CREs) [2]. Promoters and enhancers are the most common types of CREs and TF binding in these regions is ultimately responsible for activating, enhancing, and repressing gene expression programs [1,2]. Because of the role these proteins play in transcription, the discovery of TFBSs greatly furthers the understanding of many, if not all, biological processes [1]. In previous works we have used TFBS prediction to derive insights about gene expression in studies of wound healing [25], brain tumors [26], and SARS-CoV-2 [27,28].

Many tools have been created to identify TFBSs. Depending on the approach, the extent of incorporation of relevant experimental data varies widely. Early on, the position weight matrix (PWM) was used to represent and predict the binding of proteins to DNA. The PWM can then be used to obtain a likelihood score for a target DNA region, which thus represents the likelihood of a TF binding to that DNA sequence.

In the search for increased accuracy, newer models have improved TFBS prediction by incorporating other relevant biological data, such as 3D structure of DNA [3–7], chromatin accessibility/DNase hypersensitivity sites [8–10], overlap in gene ontology [11], amino acid physicochemical properties [12], and gene expression and chromatin accessibility [13,14]. These alternative models often match or outperform strictly sequence-based models [7,15] in the prediction of TFBSs, although they may involve inefficient and/or underdeveloped technologies compared with the more widely used ChIP-Seq and SELEX approaches. However, this varies by TF, and therefore, it may make sense to derive individual models composed of the most relevant contextual data for each [16]. In addition, algorithms for ab initio motif discovery and enrichment from in vivo data, such as HOMER [17], STEME [18], ProSampler [19], and STREME [20], which are reviewed here [21], are also under development. Correspondingly, several databases catalog TF motifs, most prominently JASPAR [22,23] and TRANSFAC [24].

### TFBSFootprinter incorporates multiomic transcription-relevant data

We sought to identify multiple sources of experimental data relevant to gene expression and TF binding and to incorporate them into a comprehensive model to improve the prediction of functional TFBSs. Specifically, clustering of TFBSs has been shown to be an indicator of functionality [29–31]; conservation of genetic sequences across genomes of related species is one of the most successfully used attributes in the identification of TFBSs [31,32]; proximity to the transcription start site (TSS) is strongly linked to TFBS functionality [33]; correlation of expression between a TF and another gene is an indication of a functional relationship [13,34,35]; variants in noncoding regions have a demonstrated effect on gene expression [36–38]; and variants affecting gene expression are enriched in TFBSs [39]; open chromatin regions (ascertained by ATAC-Seq or DNase sensitivity) correlate with TF binding [40]; and finally, as previously mentioned, significant effort has gone into identifying the actual composition of the binding sites themselves through the use of sequencing of TFBSs (e.g., ChIP-Seq and HT-SELEX) [41,42] as cataloged in several extensive databases, such as ReMap [43], the Cistrome Data Browser [44], and the Gene Transcription Regulation Database (GTRD) [45].

### Ensembl identifier-oriented system of analyses allows analyses in many species

For our tool, the Ensembl transcript ID was chosen as the basic unit of reference. As a result, the tool we present here — TFBSFootprinter — can offer predictions in 124 vertebrates at the time of writing, including many model organisms and domesticated and wild animals (Table 1). From human and model organisms such as mouse and zebrafish to African bush elephant, the catalog can increase as the Ensembl database itself expands. Additionally, it allows the inclusion of important datasets that are gene-centric, such as FANTOM [46] TSSs and expression data, GTEx [36] expression quantitative trait loci (eQTLs), and all annotations that are compiled within Ensembl itself. Finally, the Ensembl transcript ID provides an easy point of reference for a greater audience of scientists, thus increasing the accessibility and utility of the tool.

**Table 1.**
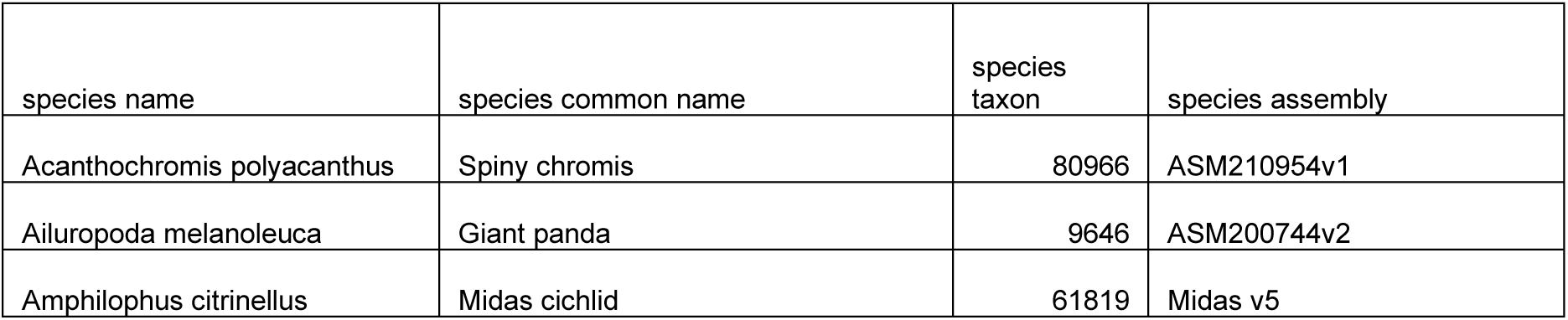
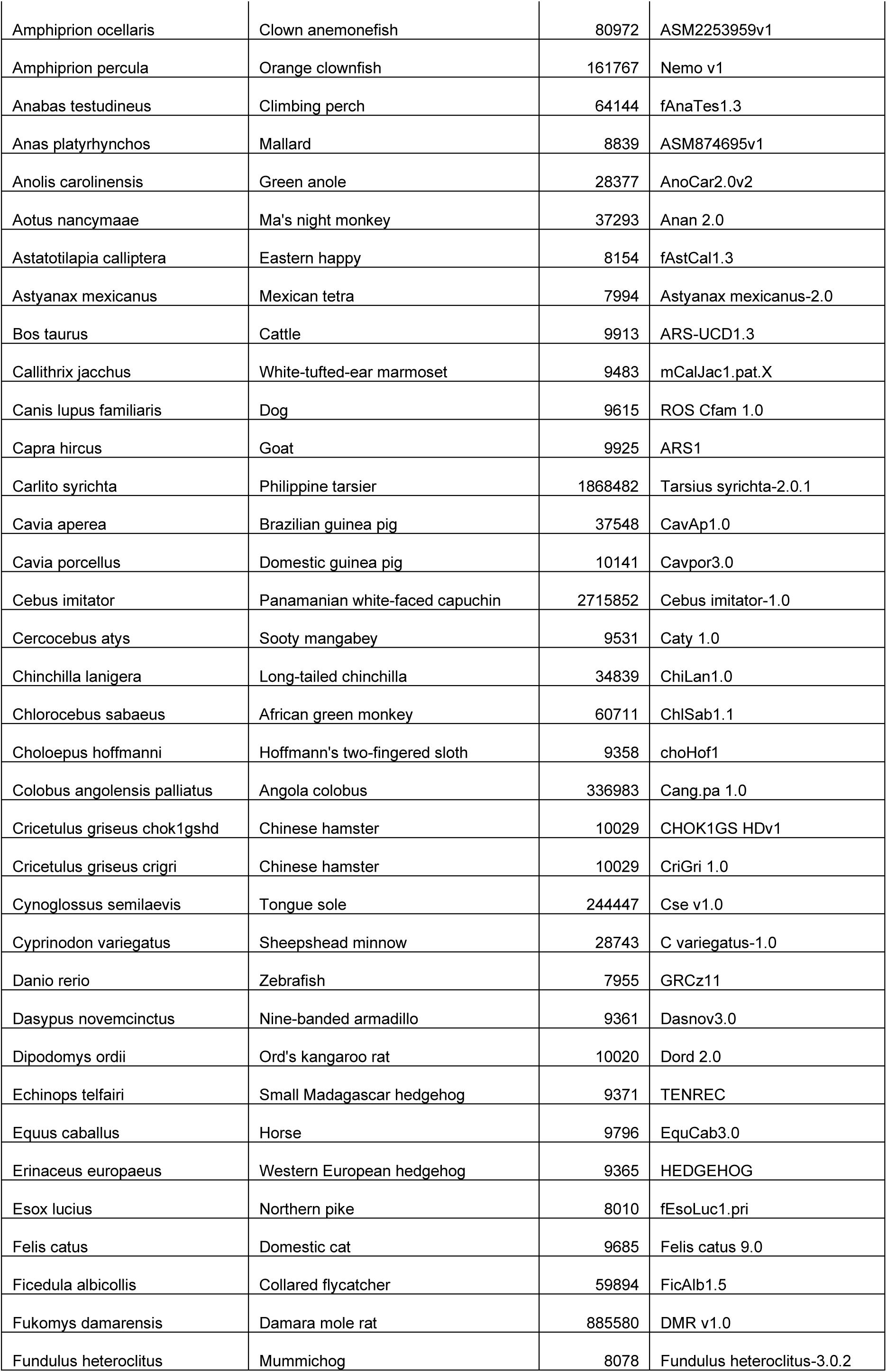
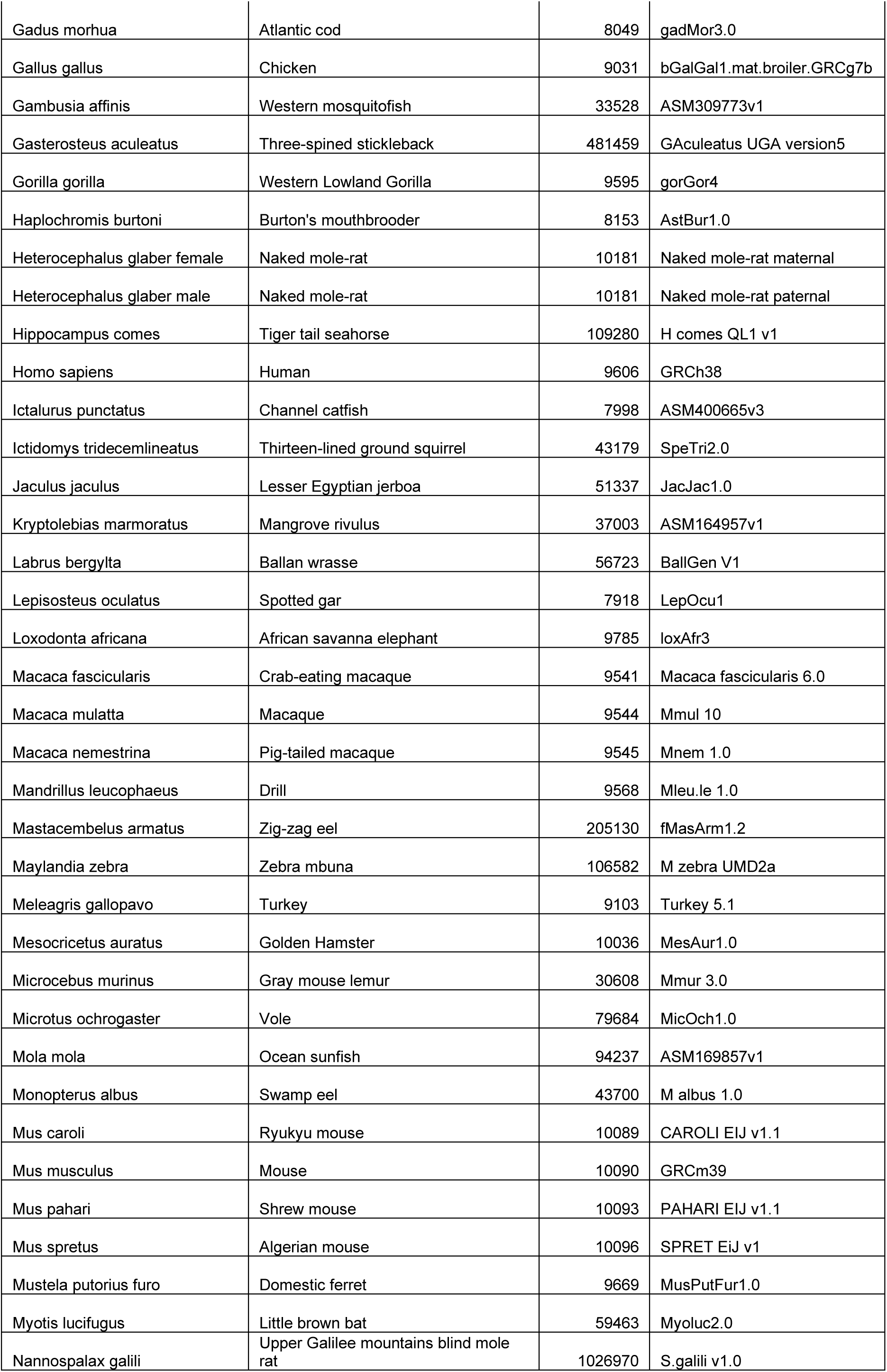
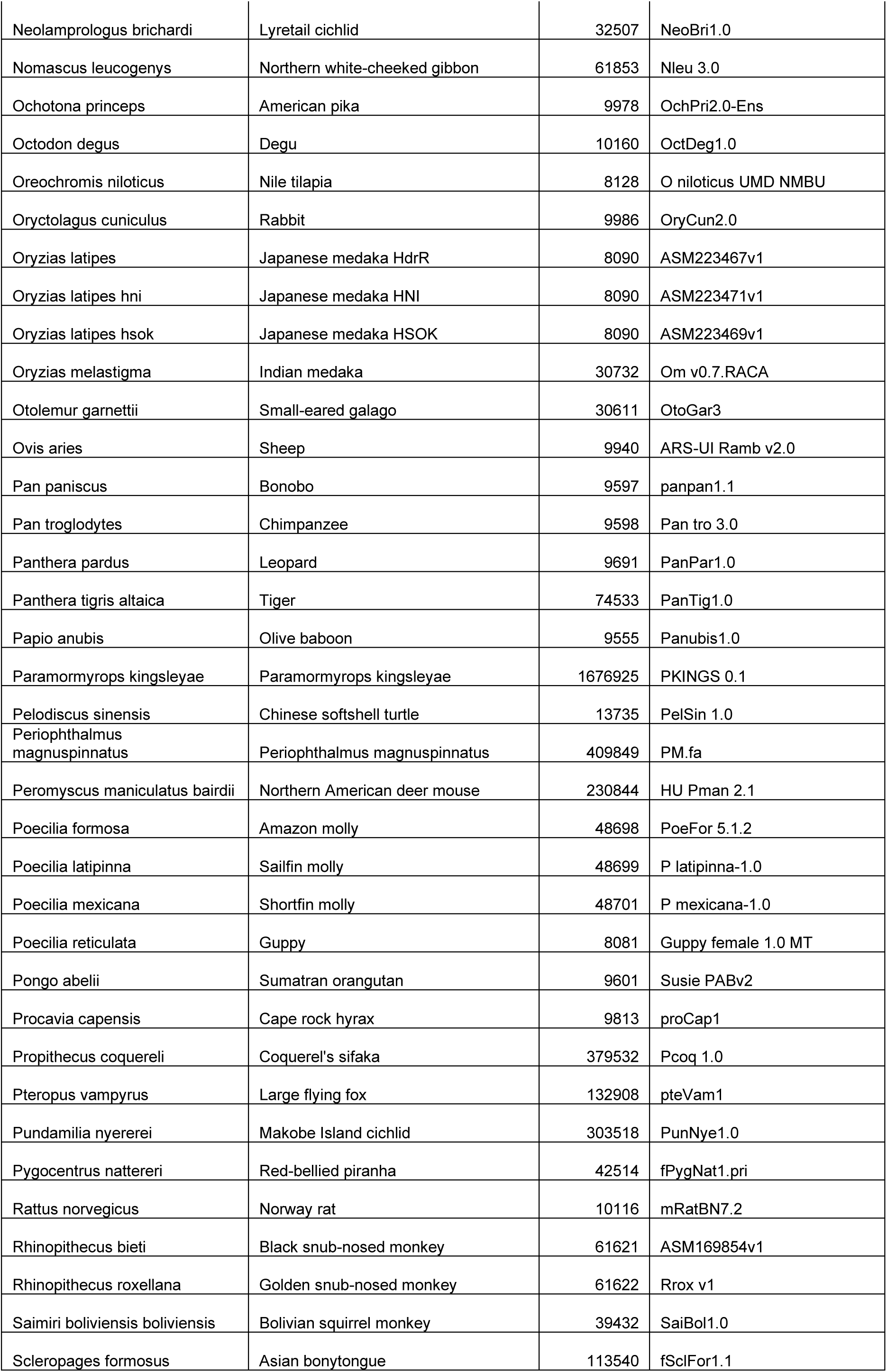
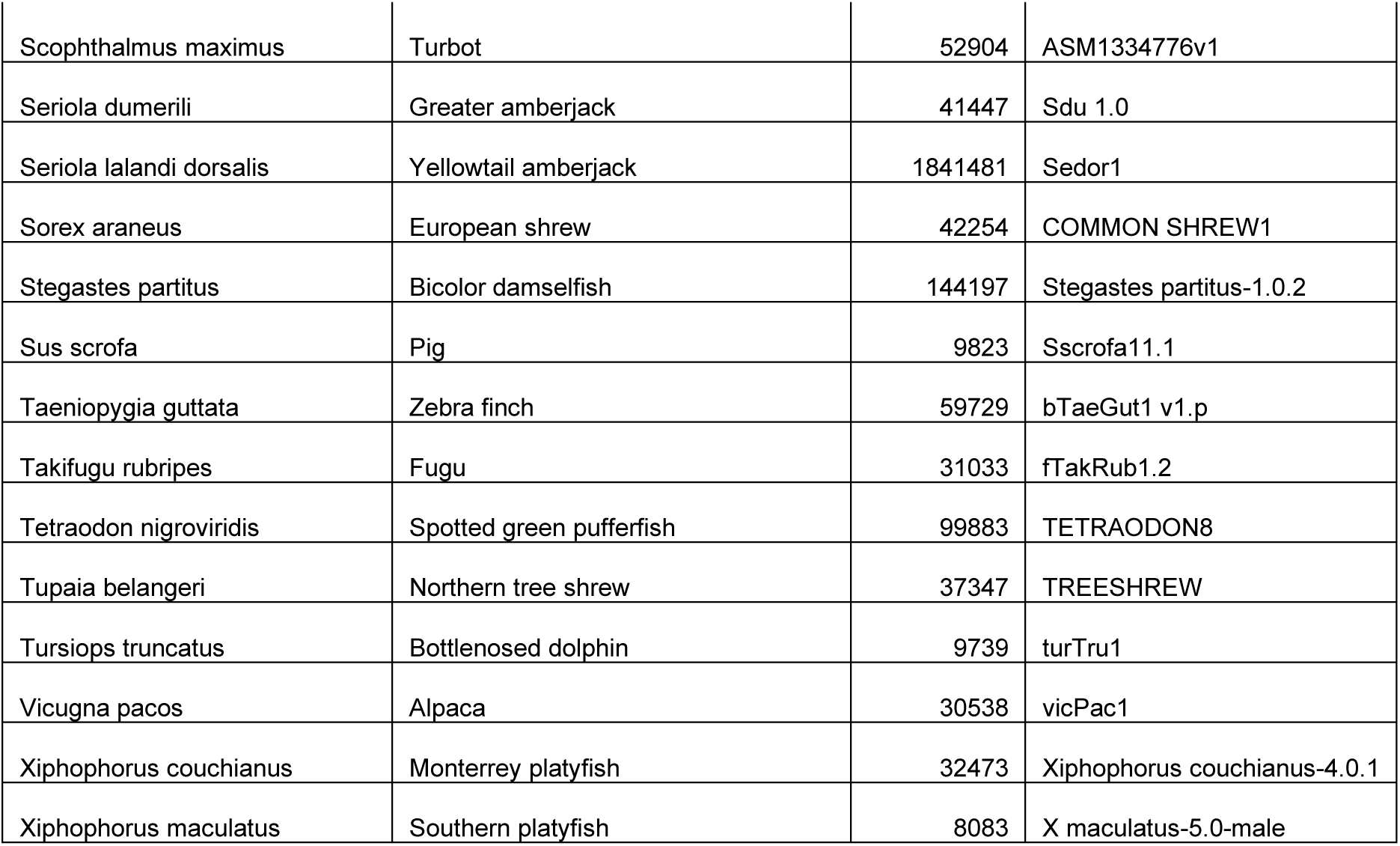
Ensembl species in which TFBSFootprinter analysis can be performed.

## Methods

### Ensembl sequence retrieval

The Ensembl Representational State Transfer (REST) server application programming interface (API) [47] is used by TFBSFootprinter for automated retrieval of user-defined DNA sequences near the transcription start site of an established Ensembl transcript ID. Annotations for the transcript and Ensembl-defined regulatory regions (e.g., ‘promoter flanking region’) are also retrieved and mapped in the final output figure.

### PWMs

A total of 575 TF position frequency matrices (PFMs) retrieved from the JASPAR database [48] (http://jaspar.genereg.net/; nonredundant) are used to create PWMs (Eq. 1), as described by [49]:

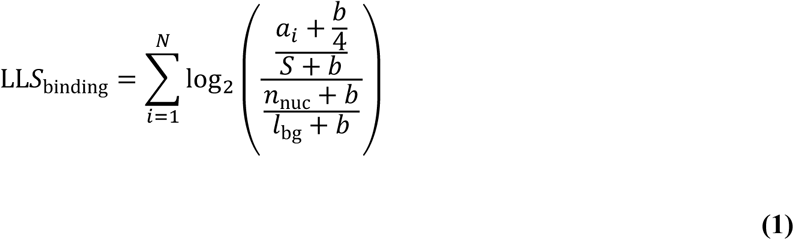

N is the set of nucleotides in the currently scanned sequence; a_i_ is the number of instances of nucleotide *a* at position i; *b* is a pseudocount set to 0.8 per [49]; *S* is the number of sequences describing the motif; n_nuc_ is the count of the nucleotide in the background sequence; and l_bg_ is the length of the background sequence. The background frequencies for each nucleotide were set to match those of the human genome as determined previously [50].

### CAGE peak locations and Spearman correlation of expression values

Cap analysis of gene expression (CAGE) uses sequencing of cDNA generated from RNA to both determine TSSs and quantify their expression levels. The FANTOM project has performed CAGE across the human genome [46], and the results are freely available for download (http://fantom.gsc.riken.jp/data/). Using the genomic locations of the FANTOM CAGE peaks, the distances from each nucleotide position in the human genome to the nearest CAGE peak were calculated. The distribution of these distances was used to generate a log-likelihood score for all observed distances. The CAGE peak locations and distance/log-likelihood score pairings are then used during de novo prediction of TFBSs (Eq. 2).

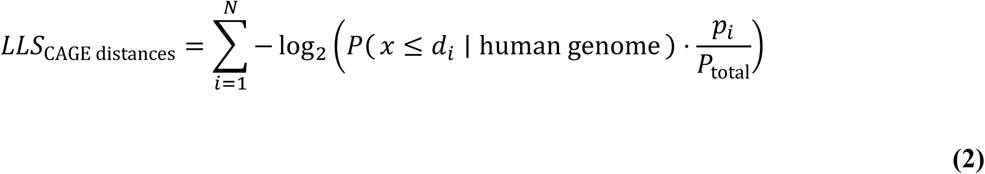

Where N is the number of all CAGE peaks associated with the target gene; d_i_ is the distance to the current CAGE peak; p_i_ is the number of peak counts of the current CAGE peak; and p_total_ is the total peak count for this gene.

The expression data for CAGE peaks associated with the 575 JASPAR TF genes were then combined with the expression data for all CAGE peaks to perform a total of 386,652,770 Spearman correlation analyses via the ‘spearmanr’ function from the SciPy Stats module [51]. Bonferroni correction was performed to account for multiple testing. Owing to the size of the analysis, a cutoff correlation magnitude value of 0.3 was used, and all the lower values (−0.3<x<0.3) and correlation pairs were discarded. A distribution was generated from the resulting correlation data, which were used to generate log-likelihood scores for each possible correlation value (Eq. 3). The CAGE peak expression correlations/log-likelihood score pairings are then used during de novo prediction of TFBSs. Computation was performed via the supercomputing resources of the CSC – IT Center for Science Ltd.

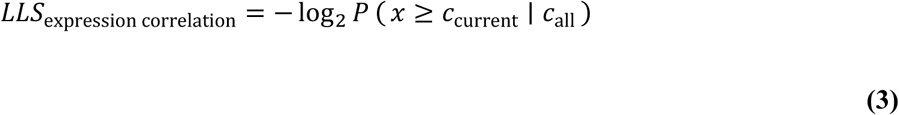

Where c_current_ is the Spearman correlation between the expression of the target gene and the expression of the TF corresponding to the putative TFBS and where c_all_ is the distribution of all Spearman correlations between JASPAR TF genes and all genes.

### Experimental TFBSs compiled by the GTRD

The GTRD project (gtrd.biouml.org) is the largest comprehensive collection of uniformly processed human and mouse ChIP-Seq peaks and has compiled data from 8,828 experiments extracted from the Gene Expression Omnibus (GEO), Sequence Read Archive (SRA), and Encyclopedia of DNA Elements (ENCODE) databases [45]. One of the outputs of the performed analyses is reads that have been grouped to identify ‘metaclusters’, places where TF binding events cluster together in the human genome. We retrieved the metacluster data (28,524,954 peaks) from the GTRD database (version 18.0) [52] and subsequently mapped the number of overlapping metaclusters for each nucleotide position in the human genome. The distribution of these overlaps was used to generate a log-likelihood score for all observed overlap counts. The metacluster locations and distance/log-likelihood score pairings are then used during de novo prediction of TFBSs (Eq. 4).

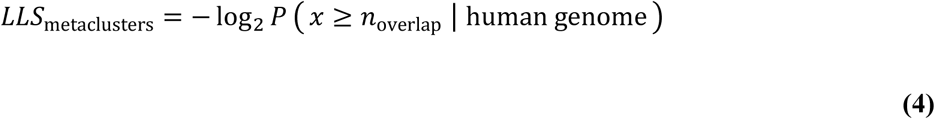

Where n_overlap_ is the number of metaclusters overlapped by the current putative TFBS and where the D_human genome_ is the distribution of the number of overlapping metaclusters for every nucleotide position in the human genome.

### ATAC-Seq peaks

The assay for transposase-accessible chromatin using sequencing (ATAC-Seq) is an experimental method for revealing the location of open chromatin [53]. These locations are indicative of genomic regions that, owing to their unpacked nature, may allow TFs to bind to DNA and subsequently influence transcription. Open chromatin regions have been shown to be useful in the prediction of TFBSs [54]. We retrieved and compiled data from 135 ATAC-Seq experiments stored in the ENCODE project database (www.encodeproject.org) and mapped the distance from each nucleotide position in the human genome to the nearest ATAC-Seq peak, and the distribution of these distances was used to generate a log-likelihood score for all observed distances. The ATAC-Seq peak locations and distance/log-likelihood score pairings are then used during de novo prediction of TFBSs (Eq. 5).

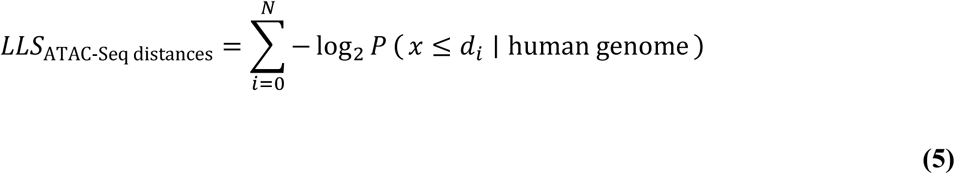

Where N is the number of ATAC-Seq peaks within the current target region; d_i_ is the distance to the current ATAC-Seq peak; and D_human genome_ is the distribution of the distances to the nearest ATAC-Seq peak for each nucleotide position in the human genome.

### eQTLs

The genome tissue expression (GTEX) project (gtexportal.org; version 7) has performed expression quantitative trait loci (eQTL) analysis on 10,294 samples from 48 tissues from 620 persons [36,55]. This analysis identified 7,621,511 variant locations in the genome, usually 1–5 base pairs (bp), that affect gene expression. eQTL data were extracted from the GTEX database and used to construct a distribution of the magnitude of effect on gene expression, which was then used to generate log-likelihood scores (Eq. 6). Next, we generated a second distribution of the distance from each gene to its variants; the distance was limited to 1,000,000 bp from either end of the transcript, as this is the search area over which GTEx scans for variants affecting the expression of each gene. The variant locations, magnitude of effect/log-likelihood score pairings, are then used during de novo prediction of TFBSs.

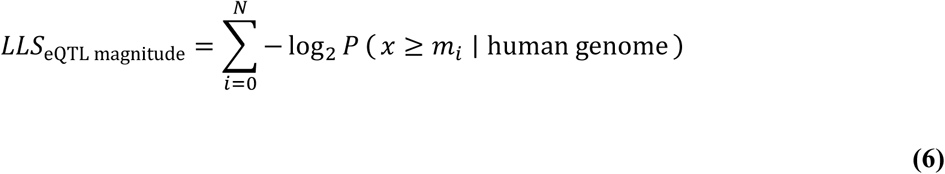

Where N is the number of eQTLs overlapping the current putative TFBS; m_i_ is the magnitude of effect of an eQTL overlapping the current putative TFBS; and D_human genome_ is the distribution of all eQTL magnitudes in each nucleotide position in the human genome.

### CpG islands

Because the methylation of DNA acts as a repressor of transcription, active promoters tend to be unmethylated. When methylated, the cytosine in a CpG dinucleotide can deaminate to thymine. Therefore, a CpG ratio close to what would be expected by chance is often indicative of an active promoter region [56,57]. Subsequently, CpG ratios (observed/expected) across a 200 nucleotide (nt) window were computed for each nucleotide position in the target genome. A distribution of these ratios was generated and used to generate log-likelihood scores for each possible ratio (Eq. 7). CpG ratio/log-likelihood score pairings are then used during de novo prediction of TFBSs.

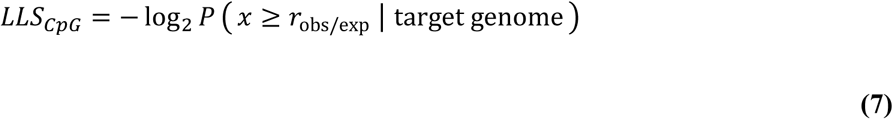

Where r_obs/exp_ is the ratio of observed to expected CpG dinucleotides in a 200 bp window centered on the current putative TFBS and where D_genome_ is the distribution of r_obs/exp_ across all nucleotide locations in the target genome.

### Conservation of vertebrate DNA

Conservation of sequence analysis has been performed by Ensembl to identify constrained elements for each species in each species group via the genomic evolutionary rate profiling (GERP) tool [58]. For each of the vertebrate species of Ensembl release 94, we calculated the distance from all nucleotides in the associated species genome to the nearest GERP constrained element and generated distributions of distances that were used to calculate log-likelihood scores for each distance (Eq. 8). GERP element distance/log-likelihood score pairings for each species are then used during de novo prediction of TFBSs in the relevant species.

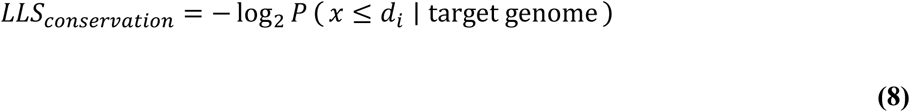

Where d_i_ is the distance between the current putative TFBS and the nearest conserved element in an alignment of 70 mammalian genomes (GERP) and D_genome_ is the distribution of distances between all nucleotides in the target genome and the nearest GERP conserved element.

### Combined affinity score

The addition of likelihood values is an established mathematical approach for measuring the combined effect of several independent parameters [59,60]. A summation of the weight (log-likelihood) scores from each experimental dataset is then performed for each putative TFBS and is represented as the ‘combined affinity score’. For analysis of human sequences, this is represented by Eq. 9. Owing to the limitations of available experimental data for nonhuman species, currently, for nonhuman vertebrates, the combined affinity score is described by Eq. 10. Complete scoring of ∼80,000+ transcript promoter regions (1,000 bp) was used to generate p values for combined affinity scoring; computation was performed via the supercomputing resources of the CSC – IT Center for Science Ltd.

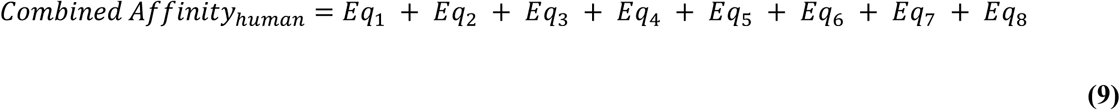

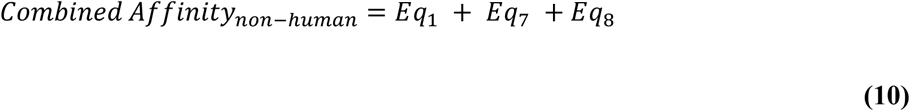

### Benchmarking of several TFBS prediction tools

For comparison, in addition to TFBSFootprinter, several other TFBS prediction tools/models were used in benchmarking: the traditional PWM, DeepBind (v. 011; SELEX and ChIP-Seq models) [61], and FIMO (meme v. 5.5.1; JASPAR 2018 nonredundant models) [62]. DeepBind was taken as an example utilizing a modern deep learning algorithm, and FIMO was chosen because it outperformed all other methods in a previous benchmark of de novo TFBS prediction tools [16]. For DeepBind, all parameters were set as defaults, analyses were performed with TF motifs on the basis of both SELEX and ChIP-Seq (when available), and the better of the two scores was retained. For TFBSFootprinter and FIMO, the p value threshold was set to 1, and all other settings were run as defaults. The TFBSFootprinter, FIMO, and PWM approaches all use JASPAR 2018 nonredundant TF motifs as the basis for scoring. For all the models, the top de novo prediction score for a target region (true positive or true negative) was kept as representative. For each TF, the correlating true positive and true negative scores were used to generate receiver operating characteristic (ROC) curves and quantify the area underneath (AUROC) via the ‘roc_curve’ module of the scikit-learn Python library [63], which is a common method for evaluating TFBS prediction [61,64–66].

### Benchmarking on experimentally verified TFBSs

Experimentally verified and curated TFBSs belonging to the annotated regulatory binding sites (ABS) [67], ORegAnno [68], and Pleiades promoter project [69] databases were retrieved as GFF files from the Pazar database [70]. From these data, 504 experimentally validated binding sites affecting gene expression for 20 DeepBind TFs and 607 experimentally validated binding sites affecting gene expression for 25 JASPAR 2018 nonredundant TFs were selected. The selection criterion for the chosen TFs was that they have at least 10 experimentally validated binding sites affecting gene expression. All target sites were converted from Hg19 to GRCh38 genomic coordinates via Ensembl REST. Subsequently, 50 bp sequences centered on each experimentally validated functional binding site in the human genome were retrieved to serve as true positives. The window length of 50 bp was chosen because it is wide enough to contain the longest TF motif (21 positions), which may overlap with the experimentally validated location at either end while also permitting some inaccuracy as to the exact center of the verified TFBS.

For each true positive, 50 true negatives were generated. True negatives were drawn at random locations within the promoter of the same Ensembl transcript of the corresponding true positive, within a 2,000 bp window (upstream and downstream) centered on each true positive, and at least 25 bp away.

### Analyzing the effect of multiomic transcription-relevant data on TFBS prediction

In addition to TFBSFootprinter benchmark scoring using all multiomic features, all 128 possible combinations of transcription-relevant features (PWM, CAGE, eQTL, metaclusters, ATAC-Seq, CpG, sequence conservation, expression correlation), which include PWM as one of the components, were used in scoring the true positives and true negatives. This allowed the identification of the best possible feature-combination TFBSFootprinter model for each TF, labeled ‘TFBSFootprinter best by TF’, as well as the TFBSFootprinter model, which performed best on average across all TFs, labeled ‘TFBSFootprinter best overall’. In the assessment of the DeepBind tool, both available models, which are based on SELEX or ChIP-Seq data, were used. Using a paired-sample t test, comparisons of ROC scores were made between all of the models: TFBSFootprinter, PWM, DeepBind, and FIMO.

## Results

### Experimental datasets used in TFBS identification

An outline of the TFBSFootprinter methodology is given in Figure 1, including the results of an example analysis of the DNA damage repair gene BRCA2 (BReast CAncer gene 2) Ensembl transcript ENST00000380152. Importantly, the results of this example prediction match previously experimentally verified TFBSs in the BRCA2 promoter, specifically sites for USF1, ELF1, and E2F family factors [71]. Several of the other predictions are for proteins that have a known role in both DNA damage, NPAS2 [72] and ID2 [73], and breast cancer [73,74].

**Figure 1.**
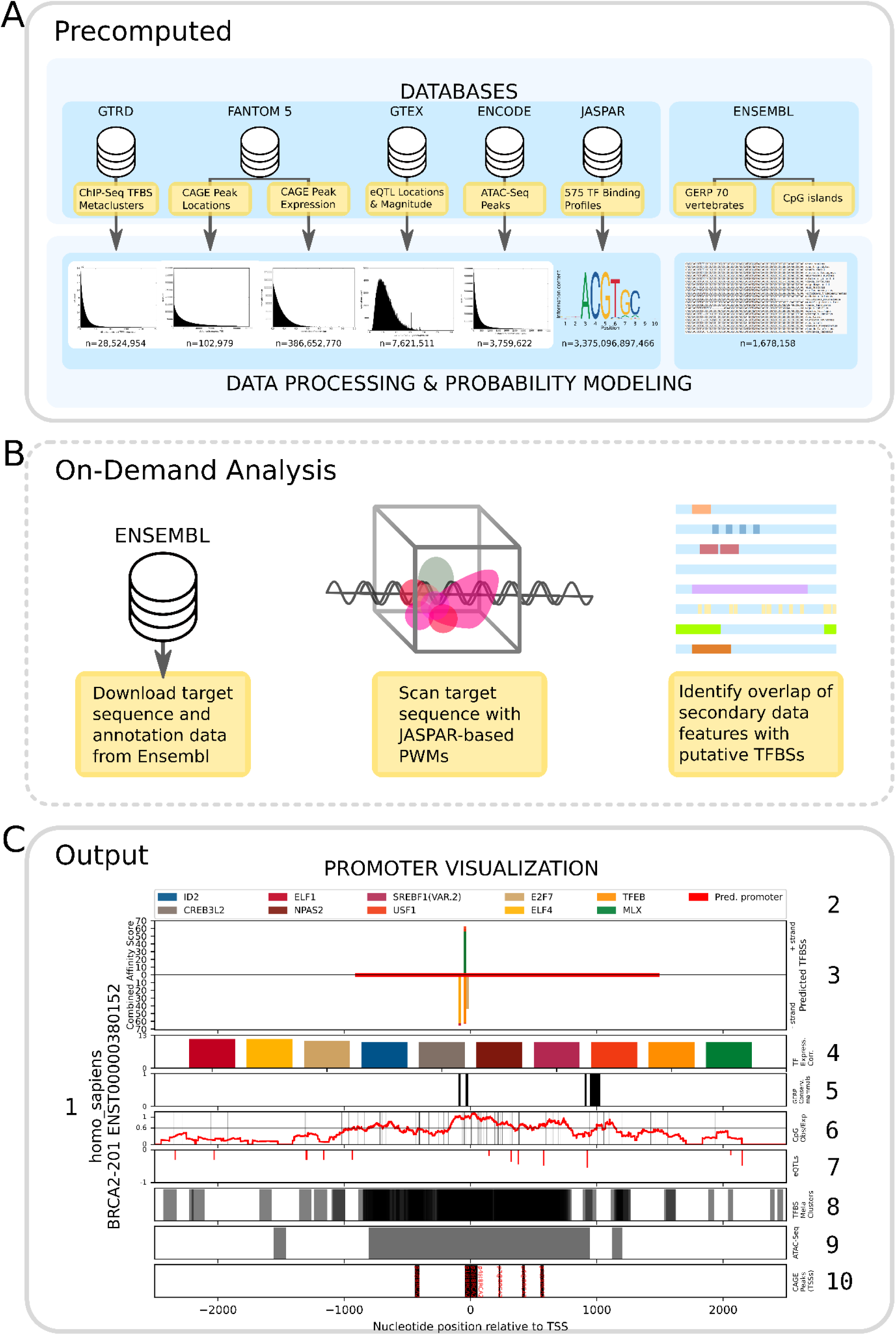
Outline of the datasets used in TFBSFootprinter. (**A**) A total of six empirical datasets are used to support the computational prediction of TFBSs via the TFBSFootprinter tool. The experimental data have been preprocessed to generate score distributions from which probability scores can be applied to putative TFBSs (n values indicate the number of elements used to compute distributions). (**B**) The user defines a target Ensembl transcript ID and TSS-related start/end sites, which are then used to download the corresponding DNA sequence and annotation data via the Ensembl API. PWM analysis of the DNA sequence generates putative TFBS hits, which are then compared with elements from the experimental datasets and scored via pregenerated log-likelihood scores relevant for the target genome. (**C**) The outputs of the TFBSFootprinter analysis are a table of results, including predicted TFBS names, locations, and scoring for each metric (not pictured), as well as individual files containing sequences and annotations. A publication-ready scalable vector graphics file (.svg) is also produced, containing several elements as indicated and described. (C1) HUGO Gene Nomenclature Committee (HGNC)-based identifier + Ensembl transcript ID. (C2) Color-coded legend of the top 10 TFs predicted to bind to this promoter. (C3) Graphical representation of the promoter of the transcript where the predicted binding sites are indicated by colored bars. The bar height indicates the combined affinity score, and the bars on the positive y-axis indicate binding on the positive (sense) strand, and the negative y-axis represents the negative (antisense) strand. (C4) Log-likelihood score of the correlation of expression between each top predicted TF gene and the target gene. (C5) Highly conserved regions of 70-mammal alignment as determined by GERP analysis (black bars). (C6) Vertical lines represent CpG locations. The red line indicates the CpG ratio of the promoter sequence over a 200 bp window. (C7) Genetic variants identified in the GTEx database that affect target gene expression (eQTLs). Green indicates a positive impact on expression (positive y-axis), and red indicates a negative impact (negative y-axis). (C8) TFBS metaclusters identified in the GTRD database (gray bars). (C9) ATAC-seq peaks (open chromatin) across many different cell types retrieved from the ENCODE database (gray bars). (C10) CAGE peaks indicating TSSs identified in the FANTOM database (black bars). The nucleotide positions at the bottom are relative to the Ensembl-defined transcription start site of the target transcript and apply to C3 and C5–C10.

Experimental data from a total of six databases were incorporated into the TFBSFootprinter algorithm for analysis of human genes (Figure 1A). Data from the relevant datasets were preprocessed to generate score distributions with which putative TFBS predictions could later be compared, as described in the Methods. Each dataset allows for scoring of transcription-relevant markers in or near putative regulatory elements identified by PWM analysis: colocalization with ChIP-Seq metaclusters; cap analysis of gene expression (CAGE) peaks, ATAC-Seq peaks, or CpG islands; correlation of expression between predicted TFs and genes of interest; colocalization of eQTLs and effects on the expression of target genes; and measurement of conservation in related vertebrate species (Figure 1C). For nonhuman vertebrates, analyses are performed on the basis of preprocessed data for PWM, CpG, and conservation. The simplicity of this piecewise approach allows for easy inclusion of additional TFBS-relevant data in the future.

### TFBSFootprinter availability

The TFBSFootprinter tool (https://github.com/thirtysix/TFBS_footprinting3) is available for installation via Conda (https://anaconda.org/thirtysix/tfbs-footprinting3) and as a Python library (https://pypi.org/project/TFBS-footprinting3/) and can subsequently be easily installed on a Linux system via the single command ‘pip install TFBS-footprinting3’. Owing to size considerations, supporting experimental data for both human and nonhuman species are downloaded on demand on first usage. Documentation on background, usage, and options is available both within the program and more extensively online (tfbs-footprinting.readthedocs.io). A listing and description of the analysis command line parameters are given in Table 2.

**Table 2.** TFBSFootprinter parameters.

| Parameter | Values | Description |
| --- | --- | --- |
| --t_ids_file, -t | [Full path to filename] | Required for running an analysis. Location of a file containing Ensembl target_species transcript ids. Input options are either a text file of Ensembl transcript ids or a .csv file with individual values set for each parameter. |
| --tf_ids_file, -tfs | [Full path to filename] | Optional: Location of a file containing a limited list of Jaspar TFs to use in scoring DNA sequence [default: all Jaspar TFs]. |
| --promoter_before_tss, -pb | 0-100,000; default, 900 | Number (integer) of nucleotides upstream of TSS to include in analysis. If this number is negative the start point will be downstream of the TSS, the end point will then need to be further downstream. |
| --promoter_after_tss, -pa | 0-100,000; default, 100 | Number (integer) of nucleotides downstream of TSS to include in analysis. If this number is negative the end point will be upstream of the TSS. The start point will then need to be further upstream. |
| --top_x_tfs, -tx | 1-20; default, 10 | Number (integer) of unique TFs to include in output .svg figure. |
| --pval, -p | 0.0000001-1; default 0.01 | P value (float) for PWM score cutoff. |
| --pvalc, -pc | 0.0000001-1; default 0.01 | P value (float) for combined affinity score score cutoff. |
| -exp_data_update, -update |  | Download the latest experimental data files for use in analysis. Will run automatically if the "data" directory does not already exist (e.g., first usage). |
| --nofig, -no |  | Do not output a figure. |

### The inclusion of empirical datasets improves TFBS prediction accuracy

The performance of both individual datasets and combinations of datasets in the identification of experimentally verified functional TFBSs was tested via receiver operating characteristic (ROC) analysis (Figure 2, Table 3). Across the 14 TFs tested with all methods, the average AUROC for were TFBSFootprinter with all features (0.881), TFBSFootprinter overall best (0.910), TFBSFootprinter best by TF (0.919); for the other models the values were, DeepBind best by TF (0.798), FIMO (0.817), and PWM (0.854).

**Figure 2.**
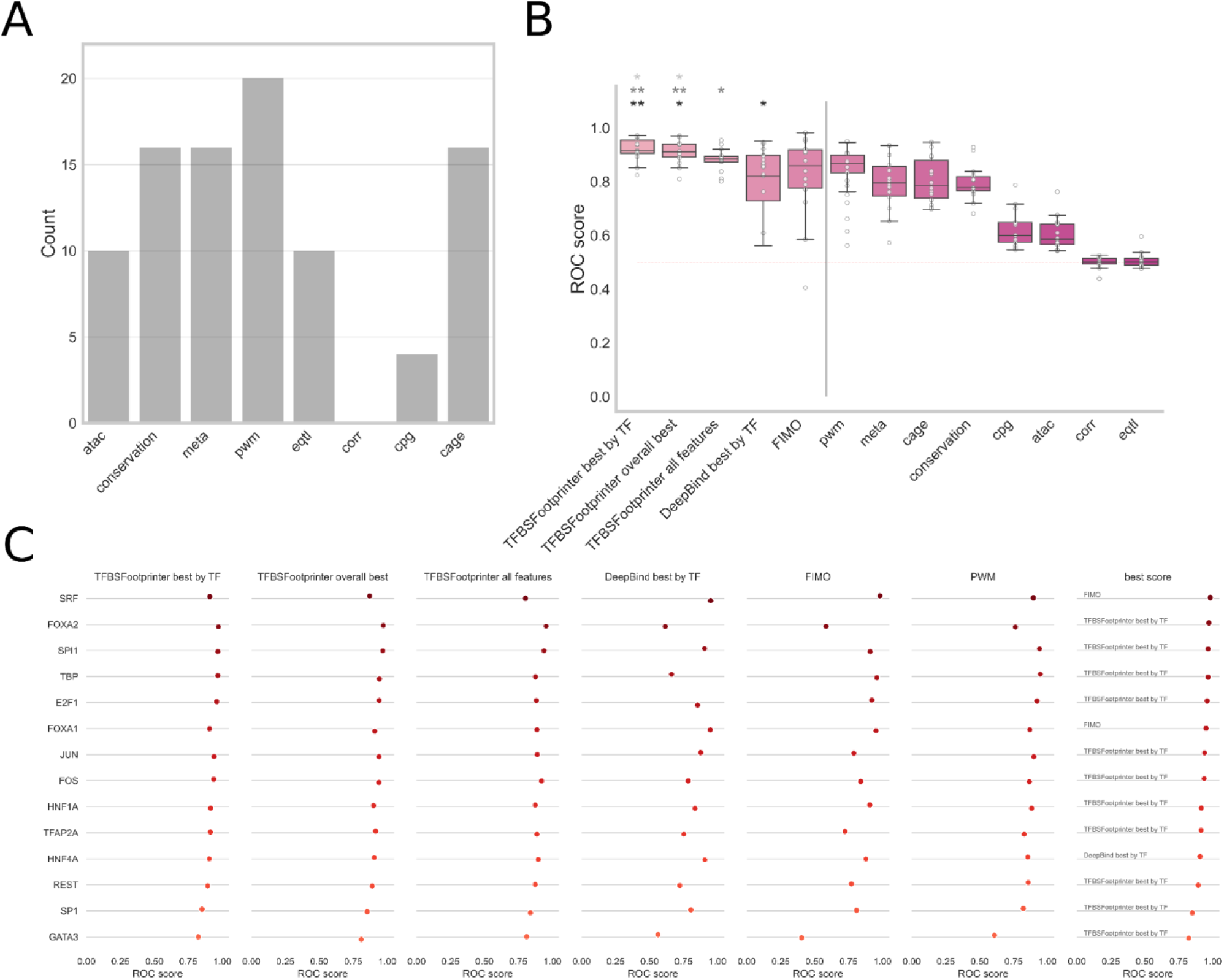
ROC analysis model performance in the identification of experimentally verified functional TFBSs—random locations in the same Ensembl transcript. ROC analysis was performed via experimentally verified functional TFBSs as annotated in the ORegAnno/Pleiades/ABS datasets as true positives, where true negatives were random locations in other Ensembl transcripts at the same distance from the TSS as the associated true positive. All the ROC curve analyses were performed on the TFs that had at least 10 true positives. and 50 true negatives per true positive were used for each analysis. Each true positive/negative segment analyzed was 50 nucleotides long, and the highest TFBS score for the relevant dataset(s) was used for each true positive/negative segment. (A) Bar plot of the frequency of experimental data types in the top 20 performing TFBSFootprinter models. (B) Boxplot of ROC scores for TFBSFootprinter, DeepBind, and FIMO for 14 TFs (left of vertical bar). ROC scores were also calculated using individual experimental metrics to show how well each contributes to accuracy of the combined model (right of vertical bar). (C) ROC scores for each individual TF tested for each primary TFBS prediction model under study. The best scoring model among all the models is named for each TF (right). TFBSFootprinter best by TF, which is based on using the highest ROC score achieved by some combination of experimental data models; TFBSFootprinter overall best, based on using the combination of experimental data models that had the best average ROC score across all the TFs analyzed; DeepBind best by TF, which is based on using the higher ROC score of the SELEX or ChIP-Seq DeepBind models.

**Table 3.**
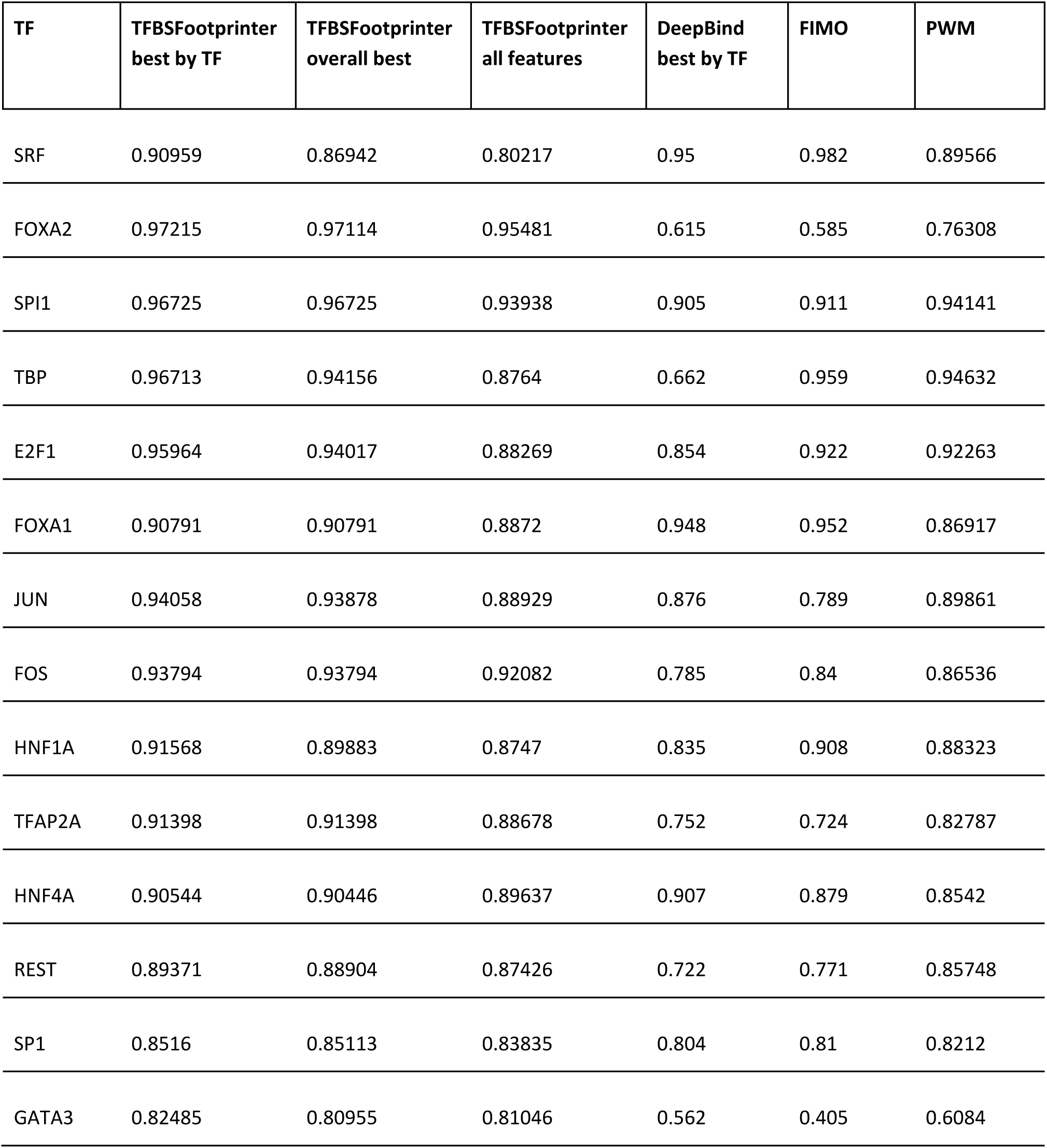
AUROC results of the TFBS prediction method.

The TFBSFootprinter model with the best average area under the ROC curve (AUROC) across all the tested TFs was the combination of PWM, ATAC, CAGE, conservation, and metacluster data (TFBSFootprinter overall best); with an average AUROC of 0.910. A paired-sample t test revealed that this model was significantly better than PWM (p value, 9.28 × 10^-3^), FIMO (p value, 3.75 × 10^-2^), and DeepBind (p value = 4.91 × 10^-3^) (Figure 2B). When all the transcription-relevant features were used, TFBSFootprinter outperformed DeepBind (p value = 3.29 × 10^-2^).

## Discussion

We have tested the newest version [25–28] of our method for the prediction of TFBSs and introduce a tool that allows analyses of promoters in 124 vertebrate species, with automatic sequence retrieval and analysis on the basis of the Ensembl transcript ID. The method leverages transcription-relevant data to augment the prediction of functional TFBSs beyond the classical PWM. In benchmarking, the TFBSFootprinter method scored evenly or better than the traditional PWM, FIMO, and DeepBind models when all the transcription-relevant data were used in its scoring. Surprisingly, benchmarking revealed that several types of transcription-relevant data, specifically eQTLs and gene expression correlations between putative TFs and target genes, did not contribute significantly to the prediction of TFBSs. As a result, the combination of features that produced the highest average ROC score across all tested TFs was PWM, ATAC, CAGE, conservation, and metaclusters. In paired-sample t test analysis, the TFBSFootprinter model performs significantly better than all the other models do. In addition, we identified specific combinations of transcription-relevant data that produced the best ROC scores for each tested TF; these combinations may lead to customizing TF models in the future. The low performance of the features related to RNA expression may be the result of the use of data that are not tissue specific; this is grounds for further research, as TFs, like many other proteins, can have tissue-specific expression patterns.

We believe that TFBSFootprinter provides an excellent way to predict TFBSs as a high-performing model overall but also because of the ease of use in working with many vertebrate organisms. TFBSFootprinter may therefore supplement current investigations into gene function or provide a means to perform larger-scale analyses of groups of related target genes. After analysis of a target transcript, a publication-ready figure depicting the top scoring TFBS candidates is produced. Additionally, a number of tables (.csv) and JavaScript Object Notation (.json) files presenting various aspects of the results are output. Primary among these is a list of computational predictions in the target species that are supported by empirical data, which are sorted by a sum of the combined log likelihood scores (the combined affinity score). Importantly, scoring of nonhuman species is limited by the availability of external data for that species; at this time, the only data commonly available for nonhuman species are sequence conservation, CpG, and JASPAR motif data. Updates of species which are available for analysis are ongoing.

### TFBSFootprinter availability

The TFBSFootprinter tool is available as a Python package and can be installed via PyPi or Conda. Any Ensembl transcript ID from any of 124 vertebrate species available in the Ensembl database can be used as input. Starting with a list of Ensembl transcript ids for a target species (e.g., Homo sapiens), TFBSFootprinter will download a user-defined region of DNA sequence from the Ensembl server. The sequence is then scored using up to 575 JASPAR TFBS profiles or a more limited set as defined by the user. Each putative TFBS is then additionally scored on the basis of transcription-relevant data, which may include, depending on the target species, proximity/overlap with the TSS, TFBS metaclusters, open chromatin, eQTLs that affect expression levels of the proximal gene, conservation of sequence, correlation of expression with the proximal (target) gene, and CpG content.

### Limitations

In the benchmark, true negatives were defined on the basis of random locations that did not overlap with experimentally verified true positives. However, there is no guarantee that these sites are indeed devoid of any binding/functionality for the TFs in question. This is a notable problem in the testing of TFBS prediction, with random sites being one of the best solutions, although it is imperfect [65]. Not all vertebrate TF binding models cataloged in the JASPAR database are applicable to every vertebrate species, as not all species possess the same genes. Users will be required to ensure that any predicted TF has an ortholog in any species in which they perform TFBS prediction.

Two of the benchmarks were scored on ChIP-Seq data from the GTRD database, and because GTRD metaclusters are part of the TFBSFootprinter scoring model, some bias is introduced. However, metaclusters are defined by merging all ChIP-Seq data (all TFs, therefore TF agnostic) across all peak-calling methods (four separate peak-calling methods), and benchmarking was performed on ChIP-Seq data from one peak-calling method for individual TFs. We chose to perform TFBSFootprinter analysis using all the available transcription-relevant features. In the future, we plan to expand the testing and assessment of empirical datasets and incorporate an option to use the combination of features that is proven best for each individual TF.

## Acknowledgements

Heini Huhtala is acknowledged for assistance in statistical techniques and professor Matti Nykter and Payam Emami Khoonsari PhD are gratefully thanked for discussions on practical and theoretical concerns. The non-profit CSC – IT Center for Science Ltd, owned by the state of Finland and Finnish higher education institutions, is acknowledged for providing computational resources for analyses.

## Data availability

The TFBSFootprinter project page is located at https://github.com/thirtysix/TFBS_footprinting3. Results of benchmarking of TFBSFootprinter (https://osf.io/hzny6/) is available as Open Science Foundation repositories.

## Abbreviations

ATAC-Seq: Assay for transposase-accessible chromatin by sequencing
bp: Base pairs
CAGE: Cap analysis of gene expression
ChIP-Seq: Chromatin immunoprecipitation with massively parallel DNA sequencing
CRE: Cis-regulatory element
eQTL: Expression quantitative trait locus
HT-SELEX: High-throughput systematic evolution of ligands by exponential enrichment
PFM: Position frequency matrix
PWM: Position weight matrix
TF: Transcription factor
TFBS: Transcription factor binding site
TSS: Transcription start site

## References

1. Lambert SA, Jolma A, Campitelli LF, Das PK, Yin Y, Albu M, et al. The Human Transcription Factors. Cell. 2018;172: 650–665. doi:10.1016/j.cell.2018.01.029

2. Wittkopp PJ, Kalay G. Cis-regulatory elements: molecular mechanisms and evolutionary processes underlying divergence. Nat Rev Genet. 2011;13: 59–69. doi:10.1038/nrg3095

3. Chiu TP, Rao S, Mann RS, Honig B, Rohs R. Genome-wide prediction of minor-groove electrostatic potential enables biophysical modeling of protein-DNA binding. Nucleic Acids Res. 2017;45: 12565– 12576. doi:10.1093/nar/gkx915

4. Rohs R, Jin X, West SM, Joshi R, Honig B, Mann RS. Origins of specificity in protein-DNA recognition. Annu Rev Biochem. 2010;79: 233–269. doi:10.1146/annurev-biochem-060408-091030

5. Zhou T, Yang L, Lu Y, Dror I, Dantas Machado AC, Ghane T, et al. DNAshape: a method for the high-throughput prediction of DNA structural features on a genomic scale. Nucleic Acids Res. 2013;41: W56–62. doi:10.1093/nar/gkt437

6. Chiu TP, Yang L, Zhou T, Main BJ, Parker SC, Nuzhdin SV, et al. GBshape: a genome browser database for DNA shape annotations. Nucleic Acids Res. 2015;43: D103–9. doi:10.1093/nar/gku977

7. Zhou T, Shen N, Yang L, Abe N, Horton J, Mann RS, et al. Quantitative modeling of transcription factor binding specificities using DNA shape. Proc Natl Acad Sci U S A. 2015;112: 4654–4659. doi:10.1073/pnas.1422023112

8. Pique-Regi R, Degner JF, Pai AA, Gaffney DJ, Gilad Y, Pritchard JK. Accurate inference of transcription factor binding from DNA sequence and chromatin accessibility data. Genome Res. 2011;21: 447–455. doi:10.1101/gr.112623.110

9. Sherwood RI, Hashimoto T, O’Donnell CW, Lewis S, Barkal AA, van Hoff JP, et al. Discovery of directional and nondirectional pioneer transcription factors by modeling DNase profile magnitude and shape. Nat Biotechnol. 2014;32: 171–178. doi:10.1038/nbt.2798

10. Deplancke B, Alpern D, Gardeux V. The Genetics of Transcription Factor DNA Binding Variation. Cell. 2016;166: 538–554. doi:10.1016/j.cell.2016.07.012

11. Qian Z, Lu L, Liu X, Cai YD, Li Y. An approach to predict transcription factor DNA binding site specificity based upon gene and transcription factor functional categorization. Bioinformatics. 2007;23: 2449–2454. doi:10.1093/bioinformatics/btm348

12. Khamis AM, Motwalli O, Oliva R, Jankovic BR, Medvedeva YA, Ashoor H, et al. A novel method for improved accuracy of transcription factor binding site prediction. Nucleic Acids Res. 2018;46: e72. doi:10.1093/nar/gky237

13. Duren Z, Chen X, Jiang R, Wang Y, Wong WH. Modeling gene regulation from paired expression and chromatin accessibility data. Proc Natl Acad Sci U S A. 2017;114: E4914–E4923. doi:10.1073/pnas.1704553114

14. Li Z, Schulz MH, Look T, Begemann M, Zenke M, Costa IG. Identification of transcription factor binding sites using ATAC-seq. Genome Biol. 2019;20: 45. doi:10.1186/s13059-019-1642-2

15. Ruan S, Stormo GD. Comparison of discriminative motif optimization using matrix and DNA shape-based models. BMC Bioinformatics. 2018;19: 86. doi:10.1186/s12859-018-2104-7

16. Jayaram N, Usvyat D, R Martin AC. Evaluating tools for transcription factor binding site prediction. BMC Bioinformatics. 2016;17: 547. doi:10.1186/s12859-016-1298-9

17. Heinz S, Benner C, Spann N, Bertolino E, Lin YC, Laslo P, et al. Simple combinations of lineage-determining transcription factors prime cis-regulatory elements required for macrophage and B cell identities. Mol Cell. 2010;38: 576–589. doi:10.1016/j.molcel.2010.05.004

18. Reid JE, Wernisch L. STEME: efficient EM to find motifs in large data sets. Nucleic Acids Res. 2011;39: e126. doi:10.1093/nar/gkr574

19. Li Y, Ni P, Zhang S, Li G, Su Z. ProSampler: an ultrafast and accurate motif finder in large ChIP-seq datasets for combinatory motif discovery. Bioinformatics. 2019;35: 4632–4639. doi:10.1093/bioinformatics/btz290

20. Bailey TL. STREME: accurate and versatile sequence motif discovery. Bioinformatics. 2021;37: 2834– 2840. doi:10.1093/bioinformatics/btab203

21. Tognon M, Giugno R, Pinello L. A survey on algorithms to characterize transcription factor binding sites. Brief Bioinform. 2023;24. doi:10.1093/bib/bbad156

22. Vlieghe D, Sandelin A, De Bleser PJ, Vleminckx K, Wasserman WW, van Roy F, et al. A new generation of JASPAR, the open-access repository for transcription factor binding site profiles. Nucleic Acids Res. 2006;34: D95–7. doi:10.1093/nar/gkj115

23. Castro-Mondragon JA, Riudavets-Puig R, Rauluseviciute I, Lemma RB, Turchi L, Blanc-Mathieu R, et al. JASPAR 2022: the 9th release of the open-access database of transcription factor binding profiles. Nucleic Acids Res. 2022;50: D165–D173. doi:10.1093/nar/gkab1113

24. Wingender E, Dietze P, Karas H, Knüppel R. TRANSFAC: a database on transcription factors and their DNA binding sites. Nucleic Acids Res. 1996;24: 238–241. doi:10.1093/nar/24.1.238

25. Barker H, Aaltonen M, Pan P, Vähätupa M, Kaipiainen P, May U, et al. Role of carbonic anhydrases in skin wound healing. Exp Mol Med. 2017. doi:10.1038/emm.2017.60

26. Karjalainen SL, Haapasalo HK, Aspatwar A, Barker H, Parkkila S, Haapasalo JA. Carbonic anhydrase related protein expression in astrocytomas and oligodendroglial tumors. BMC Cancer. 2018;18: 584. doi:10.1186/s12885-018-4493-4

27. Barker H, Parkkila S. Bioinformatic characterization of angiotensin-converting enzyme 2, the entry receptor for SARS-CoV-2. PLoS One. 2020;15: e0240647. doi:10.1371/journal.pone.0240647

28. Arppo A, Barker H, Parkkila S. Bioinformatic characterization of ENPEP, the gene encoding a potential cofactor for SARS-CoV-2 infection. PLoS One. 2024;19: e0307731. doi:10.1371/journal.pone.0307731

29. Berman BP, Nibu Y, Pfeiffer BD, Tomancak P, Celniker SE, Levine M, et al. Exploiting transcription factor binding site clustering to identify cis-regulatory modules involved in pattern formation in the Drosophila genome. Proc Natl Acad Sci U S A. 2002;99: 757–762. doi:10.1073/pnas.231608898

30. Cusanovich DA, Pavlovic B, Pritchard JK, Gilad Y. The functional consequences of variation in transcription factor binding. PLoS Genet. 2014;10: e1004226. doi:10.1371/journal.pgen.1004226

31. Hemberg M, Kreiman G. Conservation of transcription factor binding events predicts gene expression across species. Nucleic Acids Res. 2011;39: 7092–7102. doi:10.1093/nar/gkr404

32. Wenger AM, Clarke SL, Guturu H, Chen J, Schaar BT, McLean CY, et al. PRISM offers a comprehensive genomic approach to transcription factor function prediction. Genome Res. 2013;23: 889–904. doi:10.1101/gr.139071.112

33. Koudritsky M, Domany E. Positional distribution of human transcription factor binding sites. Nucleic Acids Res. 2008;36: 6795–6805. doi:10.1093/nar/gkn752

34. Haynes BC, Maier EJ, Kramer MH, Wang PI, Brown H, Brent MR. Mapping functional transcription factor networks from gene expression data. Genome Res. 2013;23: 1319–1328. doi:10.1101/gr.150904.112

35. Ma S, Snyder M, Dinesh-Kumar SP. Discovery of Novel Human Gene Regulatory Modules from Gene Co-expression and Promoter Motif Analysis. Sci Rep. 2017;7: 5557. doi:10.1038/s41598-017-05705-2

36. G. TEx Consortium. The Genotype-Tissue Expression (GTEx) project. Nat Genet. 2013;45: 580–585. doi:10.1038/ng.2653

37. Chen J, Rozowsky J, Galeev TR, Harmanci A, Kitchen R, Bedford J, et al. A uniform survey of allele-specific binding and expression over 1000-Genomes-Project individuals. Nat Commun. 2016;7: 11101. doi:10.1038/ncomms11101

38. Shi W, Fornes O, Mathelier A, Wasserman WW. Evaluating the impact of single nucleotide variants on transcription factor binding. Nucleic Acids Res. 2016;44: 10106–10116. doi:10.1093/nar/gkw691

39. Maurano MT, Humbert R, Rynes E, Thurman RE, Haugen E, Wang H, et al. Systematic localization of common disease-associated variation in regulatory DNA. Science. 2012;337: 1190–1195. doi:10.1126/science.1222794

40. Davie K, Jacobs J, Atkins M, Potier D, Christiaens V, Halder G, et al. Discovery of transcription factors and regulatory regions driving in vivo tumor development by ATAC-seq and FAIRE-seq open chromatin profiling. PLoS Genet. 2015;11: e1004994. doi:10.1371/journal.pgen.1004994

41. Jolma A, Yan J, Whitington T, Toivonen J, Nitta KR, Rastas P, et al. DNA-binding specificities of human transcription factors. Cell. 2013;152: 327–339. doi:10.1016/j.cell.2012.12.009

42. Wang J, Zhuang J, Iyer S, Lin XY, Greven MC, Kim BH, et al. Factorbook.org: a Wiki-based database for transcription factor-binding data generated by the ENCODE consortium. Nucleic Acids Res. 2013;41: D171–6. doi:10.1093/nar/gks1221

43. Hammal F, de Langen P, Bergon A, Lopez F, Ballester B. ReMap 2022: a database of Human, Mouse, Drosophila and Arabidopsis regulatory regions from an integrative analysis of DNA-binding sequencing experiments. Nucleic Acids Res. 2022;50: D316–D325. doi:10.1093/nar/gkab996

44. Zheng R, Wan C, Mei S, Qin Q, Wu Q, Sun H, et al. Cistrome Data Browser: expanded datasets and new tools for gene regulatory analysis. Nucleic Acids Res. 2019;47: D729–D735. doi:10.1093/nar/gky1094

45. Kolmykov S, Yevshin I, Kulyashov M, Sharipov R, Kondrakhin Y, Makeev VJ, et al. GTRD: an integrated view of transcription regulation. Nucleic Acids Res. 2021;49: D104–D111. doi:10.1093/nar/gkaa1057

46. Fantom Consortium, the, Riken Pmi, Clst, Forrest AR, Kawaji H, Rehli M, et al. A promoter-level mammalian expression atlas. Nature. 2014;507: 462–470. doi:10.1038/nature13182

47. Yates AD, Achuthan P, Akanni W, Allen J, Allen J, Alvarez-Jarreta J, et al. Ensembl 2020. Nucleic Acids Res. 2020;48: D682–D688. doi:10.1093/nar/gkz966

48. Khan A, Fornes O, Stigliani A, Gheorghe M, Castro-Mondragon JA, van der Lee R, et al. JASPAR 2018: update of the open-access database of transcription factor binding profiles and its web framework. Nucleic Acids Res. 2018;46: D260–D266. doi:10.1093/nar/gkx1126

49. Nishida K, Frith MC, Nakai K. Pseudocounts for transcription factor binding sites. Nucleic Acids Res. 2009;37: 939–944. doi:10.1093/nar/gkn1019

50. Yamagishi ME, Shimabukuro AI. Nucleotide frequencies in human genome and fibonacci numbers. Bull Math Biol. 2008;70: 643–653. doi:10.1007/s11538-007-9261-6

51. Virtanen P, Gommers R, Oliphant TE, Haberland M, Reddy T, Cournapeau D, et al. SciPy 1.0: fundamental algorithms for scientific computing in Python. Nat Methods. 2020;17: 261–272. doi:10.1038/s41592-019-0686-2

52. Yevshin I, Sharipov R, Valeev T, Kel A, Kolpakov F. GTRD: a database of transcription factor binding sites identified by ChIP-seq experiments. Nucleic Acids Res. 2017;45: D61–D67. doi:10.1093/nar/gkw951

53. Buenrostro JD, Giresi PG, Zaba LC, Chang HY, Greenleaf WJ. Transposition of native chromatin for fast and sensitive epigenomic profiling of open chromatin, DNA-binding proteins and nucleosome position. Nat Methods. 2013;10: 1213–1218. doi:10.1038/nmeth.2688

54. Liu S, Zibetti C, Wan J, Wang G, Blackshaw S, Qian J. Assessing the model transferability for prediction of transcription factor binding sites based on chromatin accessibility. BMC Bioinformatics. 2017;18: 355. doi:10.1186/s12859-017-1769-7

55. GTEx Consortium, Laboratory, Data Analysis &Coordinating Center (LDACC)—Analysis Working Group, Statistical Methods groups—Analysis Working Group, Enhancing GTEx (eGTEx) groups, NIH Common Fund, NIH/NCI, et al. Genetic effects on gene expression across human tissues. Nature. 2017;550: 204–213. doi:10.1038/nature24277

56. Cohen NM, Kenigsberg E, Tanay A. Primate CpG islands are maintained by heterogeneous evolutionary regimes involving minimal selection. Cell. 2011;145: 773–786. doi:10.1016/j.cell.2011.04.024

57. Long HK, Sims D, Heger A, Blackledge NP, Kutter C, Wright ML, et al. Epigenetic conservation at gene regulatory elements revealed by non-methylated DNA profiling in seven vertebrates. Elife. 2013;2: e00348. doi:10.7554/eLife.00348

58. Cooper GM, Stone EA, Asimenos G, Nisc Comparative Sequencing Program, Green ED, Batzoglou S, et al. Distribution and intensity of constraint in mammalian genomic sequence. Genome Res. 2005;15: 901–913. doi:10.1101/gr.3577405

59. Lindsay BG. Statistical Inference from Stochastic Processes. Providence, UNITED STATES: American Mathematical Society; 1988. Available: http://ebookcentral.proquest.com/lib/tampere/detail.action?docID=3112866

60. Fraser DAS, Reid N. COMBINING LIKELIHOOD AND SIGNIFICANCE FUNCTIONS. Stat Sin. 2020;30: 1–15. Available: https://www.jstor.org/stable/26892772

61. Alipanahi B, Delong A, Weirauch MT, Frey BJ. Predicting the sequence specificities of DNA- and RNA-binding proteins by deep learning. Nat Biotechnol. 2015;33: 831–838. doi:10.1038/nbt.3300

62. Grant CE, Bailey TL, Noble WS. FIMO: scanning for occurrences of a given motif. Bioinformatics. 2011;27: 1017–1018. doi:10.1093/bioinformatics/btr064

63. Pedregosa F, Varoquaux G, Gramfort A, Michel V, Thirion B, Grisel O, et al. Scikit-learn: Machine Learning in Python. J Mach Learn Res. 2011;12: 2825–2830. Available: https://www.jmlr.org/papers/v12/pedregosa11a.html

64. Mathelier A, Wasserman WW. The next generation of transcription factor binding site prediction. PLoS Comput Biol. 2013;9: e1003214. doi:10.1371/journal.pcbi.1003214

65. Sand O, Turatsinze J-V, van Helden J. Evaluating the prediction of cis-acting regulatory elements in genome sequences. In: Frishman D, Valencia A, editors. Modern Genome Annotation: The BioSapiens Network. Vienna: Springer Vienna; 2008. pp. 55–89. doi:10.1007/978-3-211-75123-7_4

66. Park S, Koh Y, Jeon H, Kim H, Yeo Y, Kang J. Enhancing the interpretability of transcription factor binding site prediction using attention mechanism. Sci Rep. 2020;10: 13413. doi:10.1038/s41598-020-70218-4

67. Blanco E, Farré D, Albà MM, Messeguer X, Guigó R. ABS: a database of Annotated regulatory Binding Sites from orthologous promoters. Nucleic Acids Res. 2006;34: D63–7. doi:10.1093/nar/gkj116

68. Lesurf R, Cotto KC, Wang G, Griffith M, Kasaian K, Jones SJM, et al. ORegAnno 3.0: a community-driven resource for curated regulatory annotation. Nucleic Acids Res. 2016;44: D126–32. doi:10.1093/nar/gkv1203

69. Portales-Casamar E, Swanson DJ, Liu L, de Leeuw CN, Banks KG, Ho Sui SJ, et al. A regulatory toolbox of MiniPromoters to drive selective expression in the brain. Proc Natl Acad Sci U S A. 2010;107: 16589–16594. doi:10.1073/pnas.1009158107

70. Portales-Casamar E, Kirov S, Lim J, Lithwick S, Swanson MI, Ticoll A, et al. PAZAR: a framework for collection and dissemination of cis-regulatory sequence annotation. Genome Biol. 2007;8: R207. doi:10.1186/gb-2007-8-10-r207

71. Davis PL, Miron A, Andersen LM, Iglehart JD, Marks JR. Isolation and initial characterization of the BRCA2 promoter. Oncogene. 1999;18: 6000–6012. doi:10.1038/sj.onc.1202990

72. Hoffman AE, Zheng T, Ba Y, Zhu Y. The circadian gene NPAS2, a putative tumor suppressor, is involved in DNA damage response. Mol Cancer Res. 2008;6: 1461–1468. doi:10.1158/1541-7786.MCR-07-2094

73. Gianni P, Matenoglou E, Geropoulos G, Agrawal N, Adnani H, Zafeiropoulos S, et al. The Fanconi anemia pathway and Breast Cancer: A comprehensive review of clinical data. Clin Breast Cancer. 2022;22: 10–25. doi:10.1016/j.clbc.2021.08.001

74. Zienolddiny S, Haugen A, Lie J-AS, Kjuus H, Anmarkrud KH, Kjærheim K. Analysis of polymorphisms in the circadian-related genes and breast cancer risk in Norwegian nurses working night shifts. Breast Cancer Res. 2013;15: R53. doi:10.1186/bcr3445

